# Novel neural signal features permit robust machine-learning of natural tactile- and proprioception-dominated dorsal column nuclei signals

**DOI:** 10.1101/831164

**Authors:** Alastair J Loutit, Jason R Potas

**Affiliations:** School of Medical Sciences, UNSW Sydney, Sydney, New South Wales, 2052, Australia; The Eccles Institute of Neuroscience, John Curtin School of Medical Research, Australian National University, Canberra, Australian Capital Territory, 2601, Australia

**Author notes:** Correspondence: Dr Jason R Potas.

**Keywords:** feature learnability, neural prosthesis, supervised back-propagation artificial neural network, brain-machine interface, cuneate, gracile

## Abstract

Neural prostheses enable users to effect movement through a variety of actuators by translating brain signals into movement control signals. However, to achieve more natural limb movements from these devices, restoration of somatosensory feedback and advances in neural decoding of motor control-related brain signals are required. We used a machine-learning approach to assess signal features for their capacity to enhance decoding performance of neural signals evoked by natural tactile and proprioceptive somatosensory stimuli, recorded from the surface of the dorsal column nuclei in urethane-anaesthetised rats. We determined signal features that are highly informative for decoding somatosensory stimuli, yet these appear underutilised in neuroprosthetic applications. We found that proprioception-dominated stimuli generalise across animals better than tactile-dominated stimuli, and we demonstrate how information that signal features contribute to neural decoding changes over a time-course of dynamic somatosensory events. These findings may improve neural decoding for various applications including novel neuroprosthetic design.

## Introduction

Neural prostheses enable users to control robotic limbs, computer cursors, or even effect movement of the users own limbs, by translating brain signals into movement control signals (Ethier *et al.*, 2012; Hochberg *et al.*, 2012; Collinger *et al.*, 2013; Gilja *et al.*, 2015; Jarosiewicz *et al.*, 2015; Bouton *et al.*, 2016; Capogrosso *et al.*, 2016; Flesher *et al.*, 2016; Ajiboye *et al.*, 2017). Currently, neural prosthetic performance is still poor compared to natural limb movements, particularly for dexterous object manipulation. However, motor control performance can be significantly improved by restoring somatosensory feedback that rapidly updates limb status, and by improved neural decoding of motor control-related brain signals.

The dorsal column nuclei (DCN) are a potential neural prosthetic target for restoring somatosensory feedback, and have recently begun to receive attention for this purpose (Richardson *et al.*, 2015, 2016; Sritharan *et al.*, 2016; Loutit *et al.*, 2017; Suresh *et al.*, 2017; Loutit *et al.*, 2019). Determining the relevance of DCN neural signal features to natural somatosensory stimuli from which they were evoked, provides insight into the neurophysiology and functional organisation of the DCN, which may inform potential neural prosthetic research in this region.

Decoding performance can be improved by increasing the amount of information acquired from neural populations by sampling from larger time windows, more neurons, or by increasing the number or quality of features used to represent neural information to a decoder. Critically, the neural population, time window sampled, and signal features used, must represent information that is highly relevant to the desired motor behaviour or presented stimulus (Wessberg *et al.*, 2000; Carmena *et al.*, 2003; Homer *et al.*, 2013; Lebedev & Nicolelis, 2017). We previously devised a metric for quantifying the relevance of neural signal features to the stimulus from which they were evoked, which we termed *feature-learnability* (Loutit *et al.*, 2019). This approach uses a simple feed-forward back-propagation supervised artificial neural network (ANN), which has several advantages over the use of state-of-the-art deep neural network (DNN) architectures (Pandarinath *et al.*, 2018): An ANN enables the quantification of information content from signal features of interest, that, for example, may be selected on the basis of: i) their neurophysiological relevance, ii) their capacity to mimic or inform electrical stimulation in sensory applications, and/or iii) are commonly used for decoding neural signals. DNNs would not permit us to directly test specific features of interest, but rather, would identify abstract features that may have little identifiable relevance. ANNs provide a relatively simple architecture which is amenable to changing the number of input features without interfering with the standardised hidden layer and learning process. However, changing the number of inputs drastically alters a DNN’s internal connectivity which is propagated throughout each subsequent layer, and thereby does not permit a standardised comparison. Finally, ANNs can perform excellent classification accuracy with significantly smaller data sets compared to DNNs.

In the present study, we used feature-learnability to assess a battery of signal features for their capacity to enhance decoding performance of DCN neural signals evoked by natural peripheral stimuli. We presented preferentially tactile or proprioceptive mechanical stimuli, whilst recording somatosensory-evoked DCN signals using a surface multielectrode array (sMEA). We extracted twenty-two features from four categories: two categories, high-frequency (HF) and low-frequency (LF), were derived from time-domain signals, and the remaining two categories were derived from frequency-domain features; high-frequency power spectral density (HF PSD), and low-frequency power spectral density (LF PSD) features. HF and LF features contain multiunit activity information. Multiunit activity has been shown in some cases to outperform spiking activity or local field potentials in offline decoding of motor tasks (Stark & Abeles, 2007) and has been successfully used to control motor neural prostheses in tetraplegic patients (Flint *et al.*, 2013; Bouton *et al.*, 2016). However, our HF and LF features include several that, to our knowledge, have not been previously used for neural decoding in this manner. The HF PSD features have been minimally investigated, but similar frequency band ranges have been used to decode motor signals in a neural prosthesis capable of effecting limb movement through neuromuscular stimulation (Bouton *et al.*, 2016). The LF PSD features have been widely investigated in motor cortex during reaching and grasping movements, or used as decoding features for electrocorticographic or electroencephalographic neural prosthetic control (Wolpaw *et al.*, 1991; Kostov & Polak, 2000; Leuthardt *et al.*, 2004; Rickert *et al.*, 2005; Flint *et al.*, 2012; Chen *et al.*, 2013; Marathe & Taylor, 2013; Bundy *et al.*, 2016).

We aimed to use feature-learnability to evaluate the above diverse features for decoding relevance to mechanically evoked tactile and proprioception-dominated stimuli. We also aimed to investigate how feature-learnability changes when the feature extraction time window is restricted, and over the time-course of stimulus presentations. We demonstrate that the HF feature category shows the best feature-learnability, and that for large time windows, only 2 HF features are required, while for shorter time windows, a more diverse feature set improves feature-learnability. Finally, we identify a highly informative feature, which, to our knowledge, has not been widely used, but may significantly improve classification of neural signals for neural prosthetic applications.

## Results

### Tactile- and proprioceptive-dominated stimuli evoked distinct patterns of neural activity

Preliminary observations of the data revealed neural activity was greatest on midline electrodes and those ipsilateral to the site of stimulus. Tactile-dominated stimuli evoked activity that was greatest at stimulus onset and/or offset. Dowel stimuli generally evoked a short and sharp burst of neural activity that peaked within 10 ms of stimulus contact/removal, whereas the brush stimuli evoked a longer, ramped burst of neural activity that was comparatively delayed at onset/offset and to reach maximum (20-30 ms). In some cases, brush stimuli evoked two initial bursts at stimulus onset. Proprioception-dominated stimuli evoked more neural activity than tactile-evoked stimuli. In general, flexion resulted in greater neural activity compared to extension, however the time to reach maximum neural activity was similar for both proprioception-dominated stimuli, which was at approximately the midpoint of the movement. Examples of filtered signals (0.55-3.3 kHz) acquired from seven electrodes in response to the four types of stimuli applied to the left forelimb are shown in Fig 1B. Features representing neural signals are shown in Fig 2.

**Fig 1.**
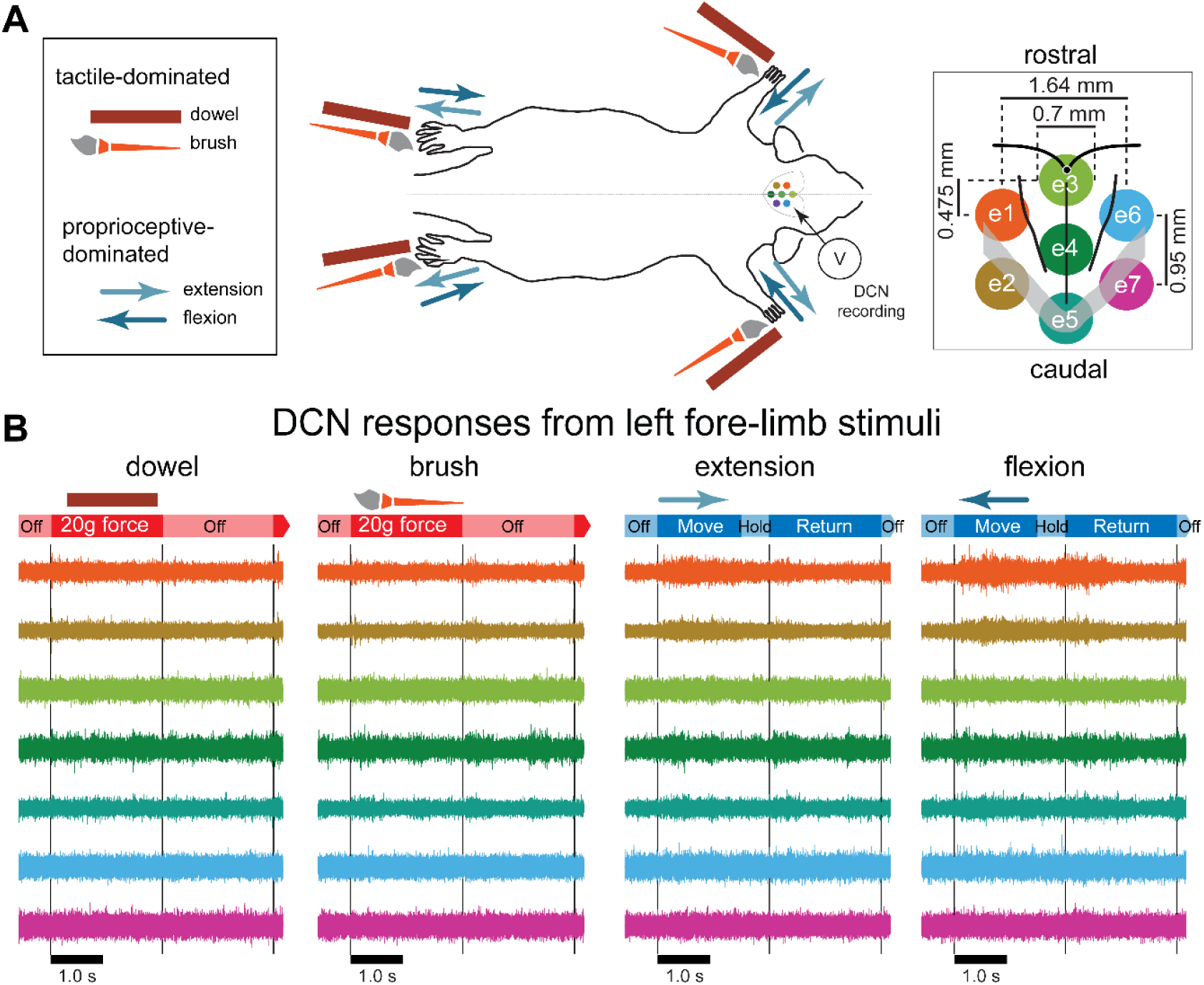
Mechanical stimulation conditions, recording arrangement, and example signals recorded from the dorsal column nuclei. **(A)** A schematic diagram of the mechanical stimulation and surface array recording paradigm. Each of the four limbs were individually stimulated with a mechanical stimulus as shown in the left insert, amounting to sixteen possible combinations of stimulus type and location. The two possible tactile-dominated stimuli comprised a phase that applied a 20-gram force with a dowel or a brush to the palmar or planter surface; the two possible proprioceptive-dominated stimuli applied extension or flexion to one of the limbs which included *move*, *hold*, and *return* phases. Dorsal column nuclei signals were simultaneously recorded from seven electrodes of a surface multi-electrode array (right insert). **(B)** Examples of 5 s of DCN signal recordings in response to stimulation of the left forelimb with each of the four stimulus types. Signals are colour-coded according to their corresponding recording electrode shown in the right insert of (A). The timing of stimulus phases is shown below each signal example for each stimulus type. Grey lines indicate the start and end of stimulus-on/-off periods. Abbreviations: DCN, dorsal column nuclei; e1, electrode 1; e2, electrode 2; e3, electrode 3; e4, electrode 4; e5, electrode 5; e6, electrode 6; e7, electrode, 7; SEM, standard error of the mean.

**Fig 2.**
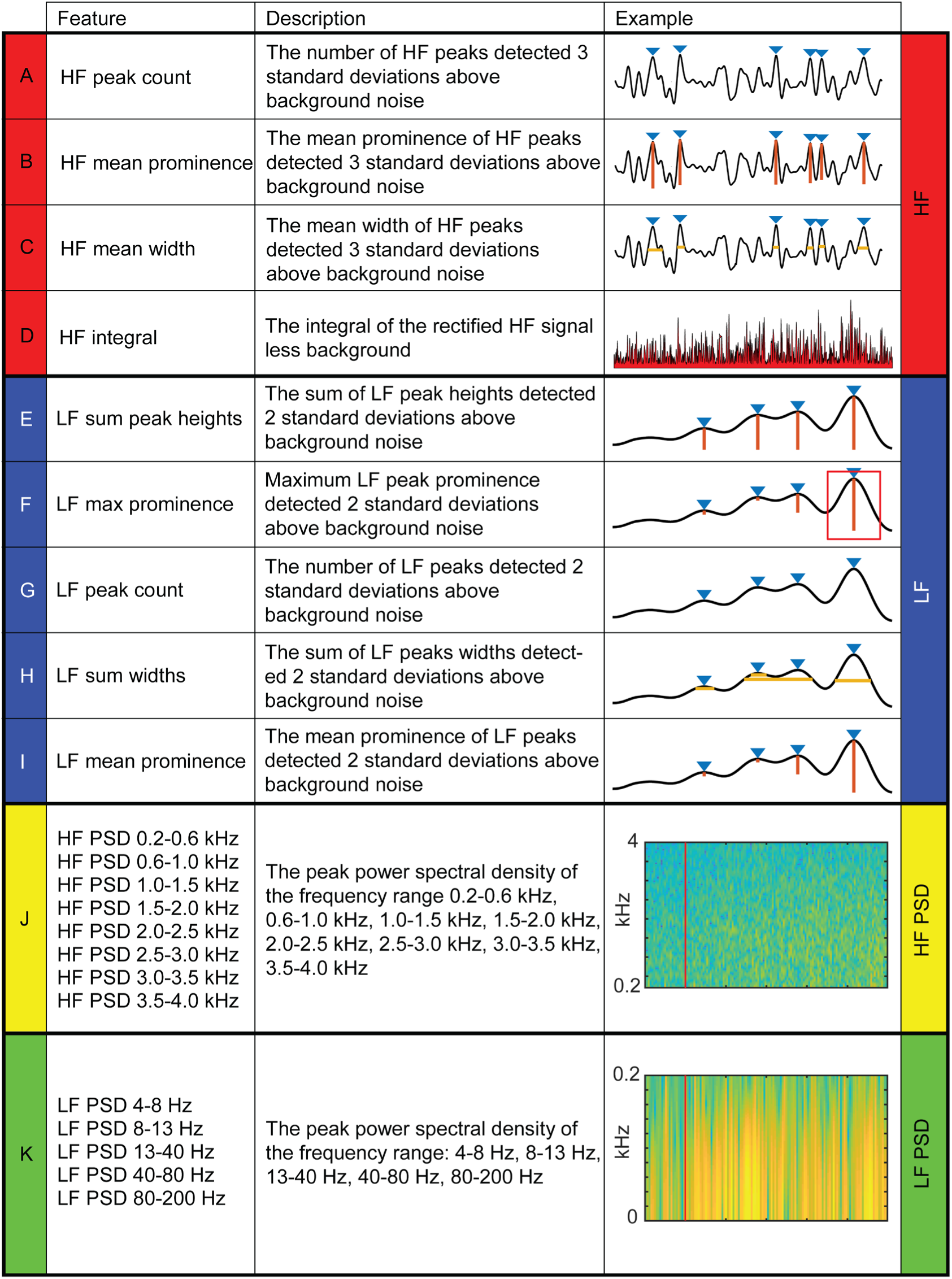
Four categories of twenty-two dorsal column nuclei signal features extracted for artificial neural network machine-learning. Descriptions and examples of individual features, extracted from dorsal column nuclei signals at various time windows, used to calculate feature-learnability (Loutit *et al.*, 2019) are shown. Individual features were divided into four categories (colour coded): four HF (**A-D**); five LF (**E-I**); eight HF PSD (**J**); and five LF PSD (**K**). Red line in J and K indicates stimulus onset. Abbreviations: HF, high-frequency, LF, low-frequency; HF PSD, high-frequency power spectral density; LF PSD, low-frequency power spectral density.

### Feature-learnability of individual features

We sought to rank individual features for their capacity to be informative of stimulus type and location. Fig 3A shows the rank order of feature-learnability for all 22 features extracted from the first 1000 ms following the stimulus onset across all 7 electrodes. All 22 features performed significantly greater than chance levels of 6.25%. Feature-learnability ranking clustered into three groups; 1) highest performing features, comprising three of the four HF and one of the five LF features; 2) middle performing features, comprising the remaining LF and all but one of the HF PSD features, and 3) the lowest performing features, comprising mainly LF PSD feature and the remaining HF and HF PSD features, all of which still performed 3 times greater than chance levels. These findings indicate that time-domain HF features are the most informative for determining a combination of stimulus location and type.

**Fig 3.**
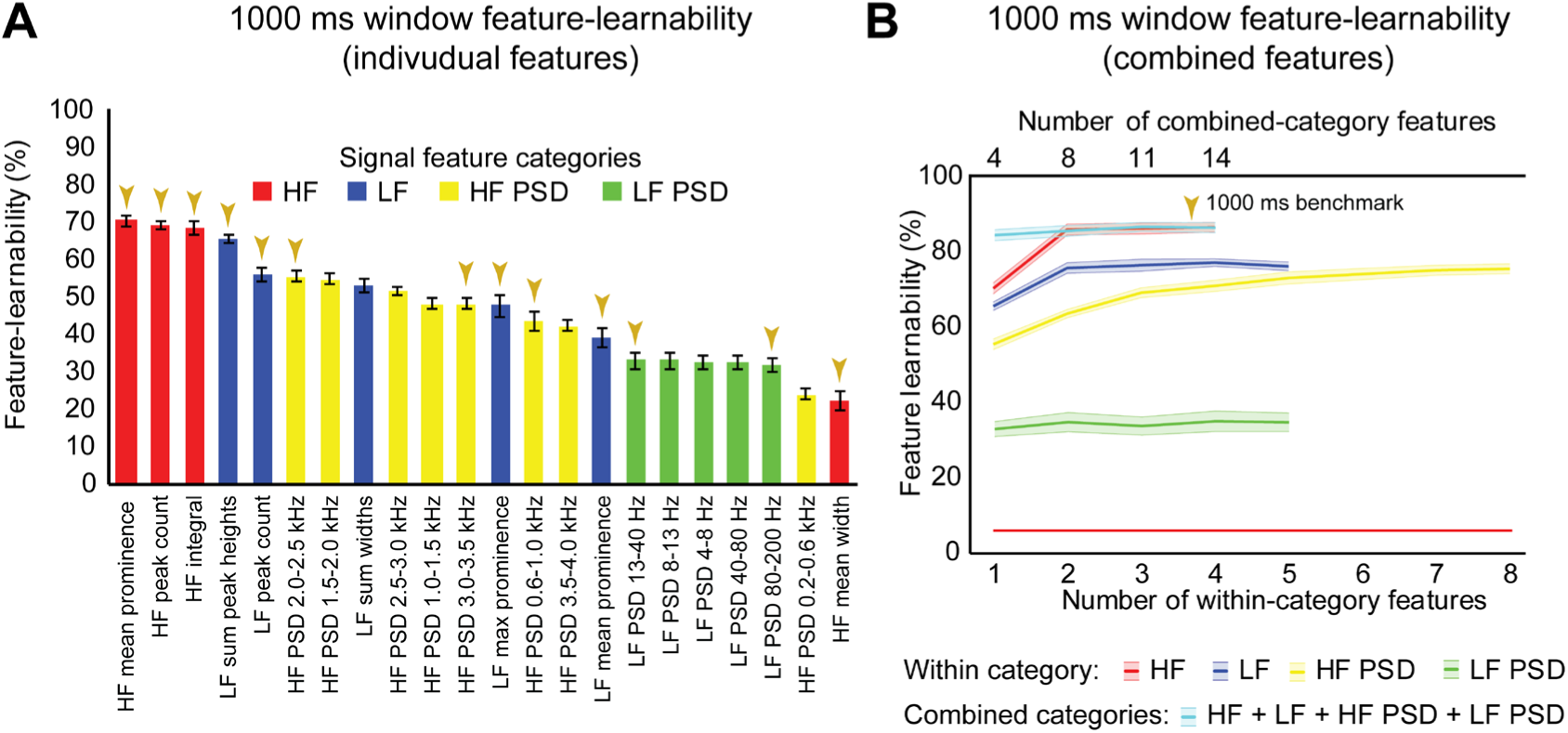
1000 ms window benchmark determined from individual and combined signal features. **(A)** Feature-learnability was derived from dorsal column nuclei signal features extracted from 1000 ms windows starting at the onset of each stimulus. Features are ordered in feature-learnability rank order (Loutit *et al.*, 2019) from highest to lowest feature-learnability from left to right. Colours indicate the feature category that individual features belong (as described in Fig 2). Gold arrows indicate the thirteen signal features that comprise the 1000 ms window benchmark configuration also indicated in B. **(B)** Feature-learnability was determined after consecutively adding individual features from the same category (within-category features) that improved classification accuracy. The number of individual features included in within-category combinations are indicated by the bottom x-axis; curves are colour coded according to the within-category features. Feature-learnability was also determined by combining the within-category features from all four categories (combined-category features, plotted in cyan). The total number of individual features included in the combined-category feature combinations is indicated by the top x-axis; the gold arrow indicates the 1000 ms window benchmark configuration for all subsequent comparisons. The red line indicates chance level of classification (6.25%). Feature learnability data expressed as mean ± SEM. Abbreviations: HF, high-frequency, LF, low-frequency; HF PSD, high-frequency power spectral density; LF PSD, low-frequency power spectral density. See Fig 2 for feature name descriptions.

### Establishing a 1000 ms window feature-learnability benchmark

We previously demonstrated that combinations of LF and HF signal features improve machine-learning outcomes (Loutit *et al.*, 2019). We therefore sought to determine a combination of input features that produced the highest feature-learnability, and to establish a benchmark for subsequent comparisons. Combinations derived from the best performing pair of features from within the HF, LF, and HF PSD categories resulted in significant feature-learnability improvements, compared to single features (p ≤ 0.01, LMER, Tukey), but tended to plateau after adding the second feature (Fig 3B). The best performing pair of frequency-domain features in the LF PSD category was not better than the best performing single LF PSD feature (p = 1.0, LMER, Tukey). The highest-ranked learnability from within-category feature combinations was achieved by combining all four HF features (86.5%). However this HF 4-feature combination was not significantly greater than the 2 or 3-feature HF combinations (p = 1.0, LMER, Tukey), all of which also significantly out-performed the best individual HF feature and all other within-category combinations (p ≤ 0.0003, LMER, Tukey; Fig 3B).

To determine if combinations of features from different categories improved feature-learnability, we combined the highest-ranked feature from each category (i.e. a combination with 4 features). This combination yielded significantly greater feature-learnability than all LF, LF PSD, and HF PSD highest-ranked feature combinations (p ≤ 0.0063, LMER, Tukey). Compared to the HF feature category, the 4-feature across-category combination was only significantly greater than the highest-ranked single HF feature (p < 0.0001, LMER, Tukey), but was not significantly different when additional HF features were added (p = 1.0, LMER, Tukey; Fig 3B).

To find the combination with the highest-ranked feature-learnability, we sequentially added the next best features in groups of four (i.e. the next best individual feature from each of the four categories). The highest-ranked feature-learnability achieved was 87.2± 1.3% with 13 features which we defined as our 1000 ms window feature-learnability benchmark for subsequent comparisons. Despite improved feature-learnability with 13 features compared to 4, 8, or 11 features, there were no significant differences between feature-learnability outcomes of any of the across-category combined feature sets (p ≥ 0.66, LMER, Tukey; Fig 3B). The individual features that contribute to the 1000 ms benchmark combination are indicated in Fig 3A, and their confusion matrices are shown in Fig 4A-D.

**Fig 4.**
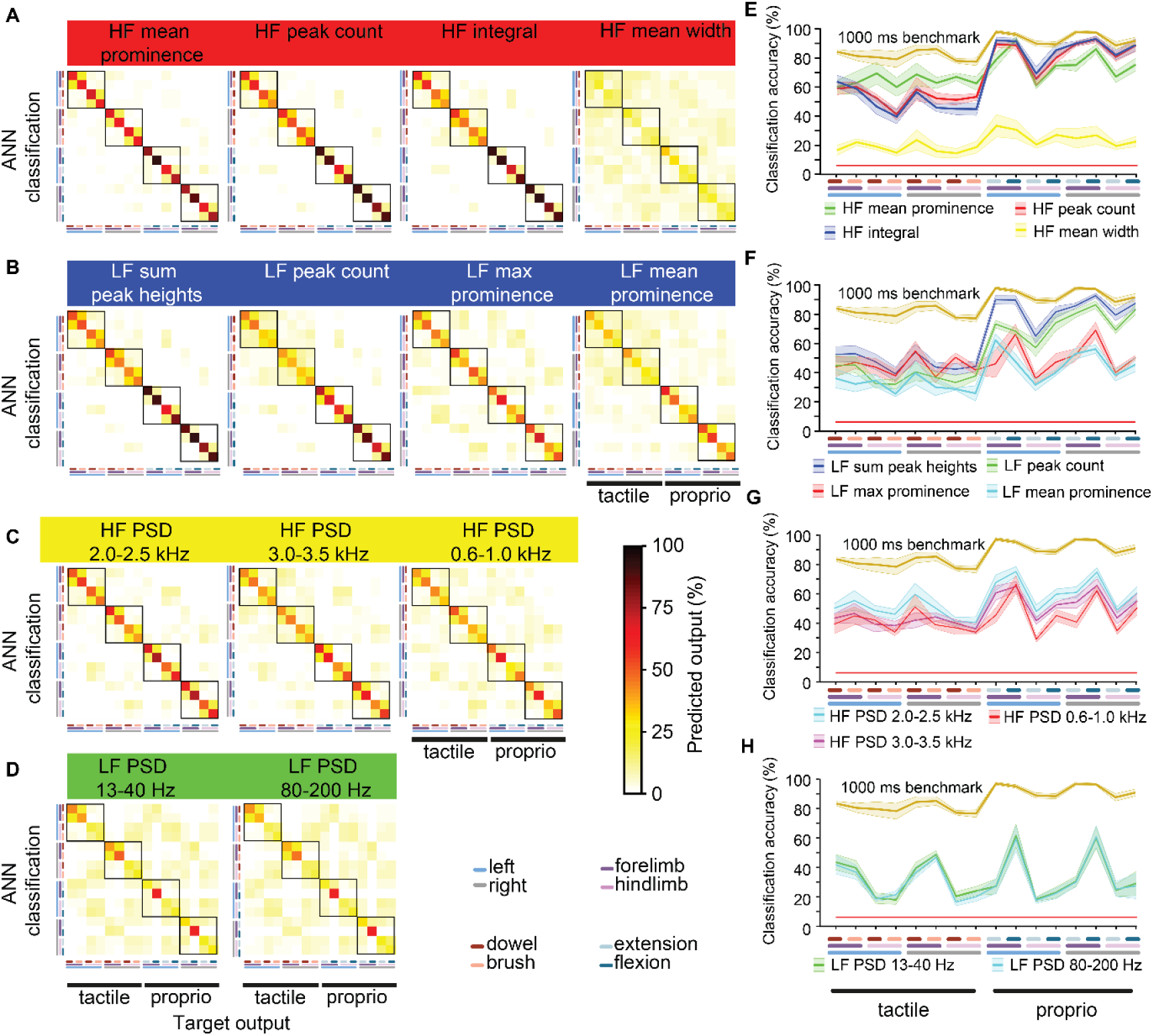
Machine-learning outcomes for individual and combinations of 1000 ms window benchmark features. **(A-D)** Shown are confusion matrices (mean of 6 animals) of machine-learning outcomes from thirteen features comprising the benchmark feature set (Fig 3A) grouped by their four categories (Fig 2). Features in the left column were the first four features combined (blue curve, Fig 3B), and each confusion matrix to the right shows successive additions to the feature set. **(E-H)** The diagonal of each confusion matrix is plotted in the line graphs, with features in (A-D) corresponding to line graphs (E-H), respectively. Dark lines in (E-H) show means of six animals and the corresponding pale bars indicate ± SEM; colours arbitrarily chosen. Superimposed is the benchmark feature learnability (gold) for comparison. See Fig 2 for feature descriptions.

In summary, time-domain HF features resulted in the highest feature-learnability rankings, and although the benchmark feature set included thirteen features from all four categories, benchmark feature-learnability was not significantly higher than the combination of two HF features (*HF mean prominence* and *HF peak count*).

### How well do individual features predict different mechanical somatosensory stimuli?

To determine what information the 1000 ms benchmark input features contribute to feature-learnability, we plotted confusion matrices for all thirteen features (Fig 4). Correct predictions (i.e. the diagonal) of each matrix are replotted with their SEM in the right panels of Fig 4E-H to facilitate performance comparisons of each feature and to provide a measure of variability among animals.

The HF feature category (Fig 4A) significantly outperformed all other categories (p ≤ 0.001, LMER, Tukey). Proprioception-dominated stimuli were significantly better classified than tactile-dominated stimuli by all categories (p < 0.0001, LMER, Tukey), except the LF PSD category (Fig 4H, p = 0.74, LMER, Tukey). The forelimbs were significantly better classified than hindlimbs across all categories (p ≤ 0.004, LMER, Tukey), and for both tactile- and proprioceptive-dominated stimuli (p < 0.0001, LMER, Tukey). *HF mean prominence* was the best predictor of tactile-dominated stimuli and significantly outperformed all other features (p ≤ 0.0026, LMER, Tukey; Fig 4E-H). *HF integral* was the best predictor of proprioceptive-dominated stimuli and significantly outperformed most other features (p ≤ 0.0019, LMER, Tukey), except *HF peak count* and *LF sum peak height* (p 1.0, LMER, Tukey) (Fig 4E-H). Interestingly, *LF peak count* predicted proprioceptive-dominated stimuli significantly better when evoked from right limbs compared to left limbs (p = 0.035, LMER, Tukey; Fig 4F).

### HF feature quantification

To investigate how *HF mean prominence* predicted tactile stimuli significantly better than all other features, including the highest performing feature *HF peak count*, we quantified *HF mean prominence* and compared this to *HF peak count* acquired from each electrode and animal. We previously demonstrated that a feature’s learnability is correlated to the number of instances different stimuli evoke significantly different magnitudes of that feature (Loutit *et al.*, 2019). We therefore quantified the two features from anatomically relevant electrodes, i.e. features acquired from stimuli applied to left forelimbs were quantified from left electrodes (e1 and e2), right forelimbs from right electrodes (e6 and e7), and hindlimbs from midline electrodes (e3, e4, e5).

Although the effect size was small, *HF mean prominence* evoked by dowel stimuli (30.2 ± 1.2 µV) were significantly higher than brush stimuli (29.7 ± 1.0 µV; p = 1.3e-4, paired *t*-test), as were *HF peak counts* (dowel, 41.6 ± 2.5 events; brush, 40.0 ± 1.9 events; p = 0.041, paired *t*-test). *HF mean prominences* evoked by flexion (32.4 ± 1.2 µV) were significantly higher than when evoked by extension (31.1 ± 1.2 µV; p = 2.1e-12, paired *t*-test), as were *HF peak counts* (flexion, 129.1 ± 8.2 events; extension 87.9 ± 5.9 events; p = 3.3e-14). Moreover, *HF mean prominences* of proprioception-dominated stimuli (31.7 ± 0.5 µV) were significantly higher than when evoked from tactile-dominated stimuli (29.8 ± 0.5 µV; p = 0.013, Student’s *t*-test), and proprioception-evoked *HF peak counts* (108.5 ± 5.7 events) were more than 2.5 times larger than when tactile-evoked (40.8±2.0 events; p = 8.4e-24, Student’s *t*-test).

In summary, despite small effect sizes, the difference in the *HF mean prominence* evoked from dowel vs. brush stimuli, resulted in a much smaller probability value compared to *HF peak counts*, which may account for the improved tactile classification outcome from *HF mean prominence*. Dowel, flexion, and proprioception-dominated stimuli evoke higher *HF mean prominence* and *HF peak counts* than brush, extension, and tactile-dominated stimuli, respectively.

### Signal feature robustness across animals

How generalisable are signal features of the 1000 ms benchmark across different animals? To answer this question, we used the LOO approach to measure how features extracted from an individual animal perform when presented to a neural network trained by the same features derived from the remaining cohort of animals. This approach measures how features from one animal generalise to all other animals (Loutit *et al.*, 2019). Compared to the WIA approach derived from neural networks optimised for individual animals using 13 features over 1000 ms (Fig 5A), feature-learnability determined by the LOO approach (Fig 5B) was significantly reduced, by almost half to that of the WIA approach (Fig 5C; p < 2.2e-16, LMER). More than 50% of the reduction in feature-learnability derived by the LOO approach was accounted for by greater confusion errors associated with tactile-dominated stimuli, specifically that dowel and brush stimuli were generally poorly discriminated, which were exacerbated by left/right errors for the hindlimb (Fig 5B-C). Compared to the WIA approach, proprioception-dominated stimuli from the LOO approach demonstrated an insignificant 11% reduction in classification accuracy at the forelimb (p = 0.25, LMER, Tukey), but a 43% reduction associated with the hindlimb (p < 0.0001, LMER, Tukey) (Fig 5C). These results indicated that the thirteen features contain information that is highly generalisable among animals for proprioceptive-dominated stimuli of the forelimb, but much less so for hindlimb proprioception and forelimb and hindlimb tactile-dominated stimuli.

**Fig 5.**
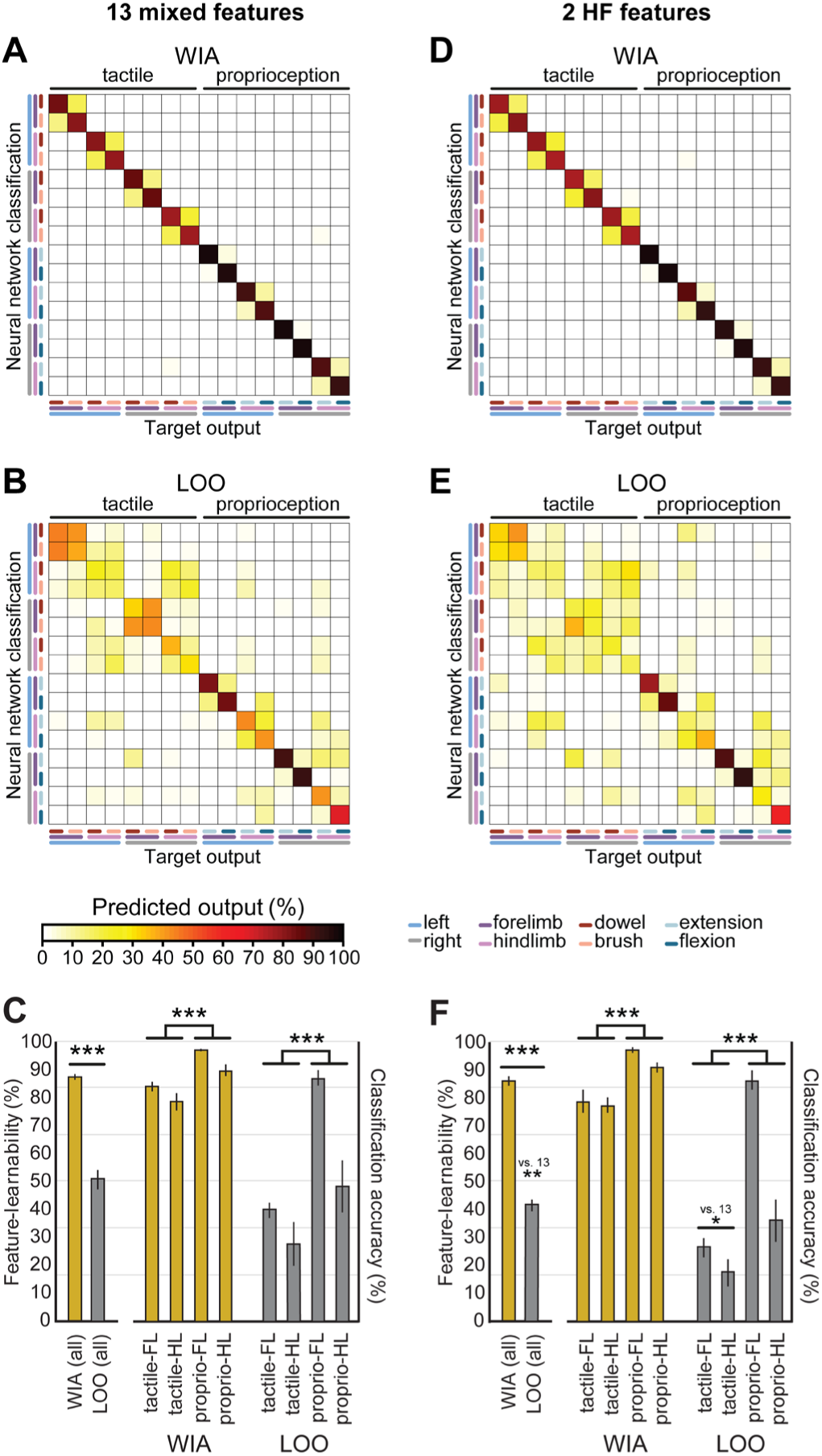
Feature robustness across animals. Confusion matrix means are shown for WIA and LOO machine-learning models. These models use identical neural network architecture extracted from 1000 ms post-stimulus onset, but differ by their training, validation, and testing data partitioning approaches. **(A)** The confusion matrix shows the mean values derived using the WIA approach for all six individual animals using the 13 mixed features that constitute the 1000 ms feature-learnability benchmark (Fig 3). The WIA approach partitions training, validation, and testing datasets from within individual animals (Loutit *et al.*, 2019), thus machine-learning models are optimised for each animal. **(B)** The confusion matrix shows the mean values derived using the LOO approach for all six individual animals on the same input data set as A. The LOO approach trains machine-learning models from all other (n = 5) animals and tests on the remaining animal, and thereby quantifies feature-learnability for each animal against the background of all others. **(C)** A comparison of the feature-learnability (left bars) for the WIA (gold bars) and LOO (grey bars) approaches are shown (derived from the means ± SEM of the diagonals in A and B respectively). The large significant reduction of feature-learnability indicates that a large portion of information encoded in the signal features were unique to individual animals, but that a significant portion also generalises across animals. The right bars show classification accuracy for stimulus classes across limbs (derived from means ± SEM calculated from the 6 confusion matrices from all animals) for WIA (gold) and LOO (grey) approaches. LOO output demonstrates the 13 signal features are highly informative for all animals for the forelimb proprioception class, but significantly less informative for all hindlimb and tactile classes. **(D)** Same analysis as A, but WIA inputs restricted to the two best HF features (*HF mean prominence* and *HF peak count*) (Fig 3). **(E)** Same data set as D using the LOO approach. **(F)** Identical analysis as per C but for the two best features (D-E). The LOO approach output for these two best features demonstrates a similar pattern shown for the 13 best features shown in C, indicating that *HF mean prominence* and *HF peak count* generalise across animals for decoding forelimb proprioception-dominated stimuli. Abbreviations: WIA, within individual animals; LOO, leave-one-out; LF, forelimb; HF, hindlimb.

Combining the two highest-ranked HF features (Fig 3) resulted in WIA feature-learnability (85.6 ± 1.2, Fig 5D) not significantly different to the combination of thirteen features (87.2 ± 1.3, Fig 5A, p = 0.13, LMER). To determine how these two features alone generalise across animals, the LOO approach was applied (Fig 5E-F). The LOO approach restricted to the two highest-ranked HF features revealed an almost identical pattern as the 13 features data set, but with the addition of forelimb/hindlimb confusion for tactile-dominated stimuli (Fig 5E-F). Compared to the 13-feature combination, LOO feature-learnability of the 2-feature combination was overall significantly reduced (p = 0.002, LMER, Fig 5F), which resulted because of reduced tactile stimulus classification accuracy that was generalised across both hind- and forelimbs (p = 0.025, Tukey, Fig 5E-F). Interestingly, there was no significant reduction in proprioception performance in the LOO approach between the 13- and 2-feature sets (p = 0.37, LMER, Tukey).

These results indicate that the two HF features extracted over 1000 ms can provide almost all of the information provided by the larger 13-feature set for forelimb proprioceptive-dominated stimulation, but the other 11 features contribute some additional information, common across all animals, which is important to discriminate hindlimb proprioceptive-stimuli as well as fore-and hindlimb tactile-dominated stimuli.

### Learnability of features extracted over different time windows

Features extracted over a 1000 ms period have little practical value for neural prosthetic feedback control. A DCN-targeted neural prosthetic device would need to provide stimulus features over shorter time windows to enable rapid updating of limb sensory status. To determine how learnable DCN signal features are over shorter periods, feature-learnability was calculated for each signal feature extracted over time windows ranging from 20-500 ms starting from stimulus onset (Fig 6A-D). A window of 60 ms was determined as the time of most abrupt change in feature-learnability across all features (see arrows, Fig 6A-D) before reaching a plateau, where feature-learnability improved slightly, or not at all, until much larger time windows (≥ 250 ms). Feature-learnability rank order derived from features extracted from 60 ms windows (Fig 6E) was altered compared to 1000 ms windows (Fig 3A). Notably, *HF integral* out-ranked *HF mean prominence* (the best feature from 1000 ms windows), although these features were not significantly different (p = 0.052, LMER, Tukey; Fig 6E), and the second-ranked feature was *LF sum peak heights* (ranked 4^th^ from 1000 ms windows). *HF mean width* remained the lowest ranked feature.

**Fig 6.**
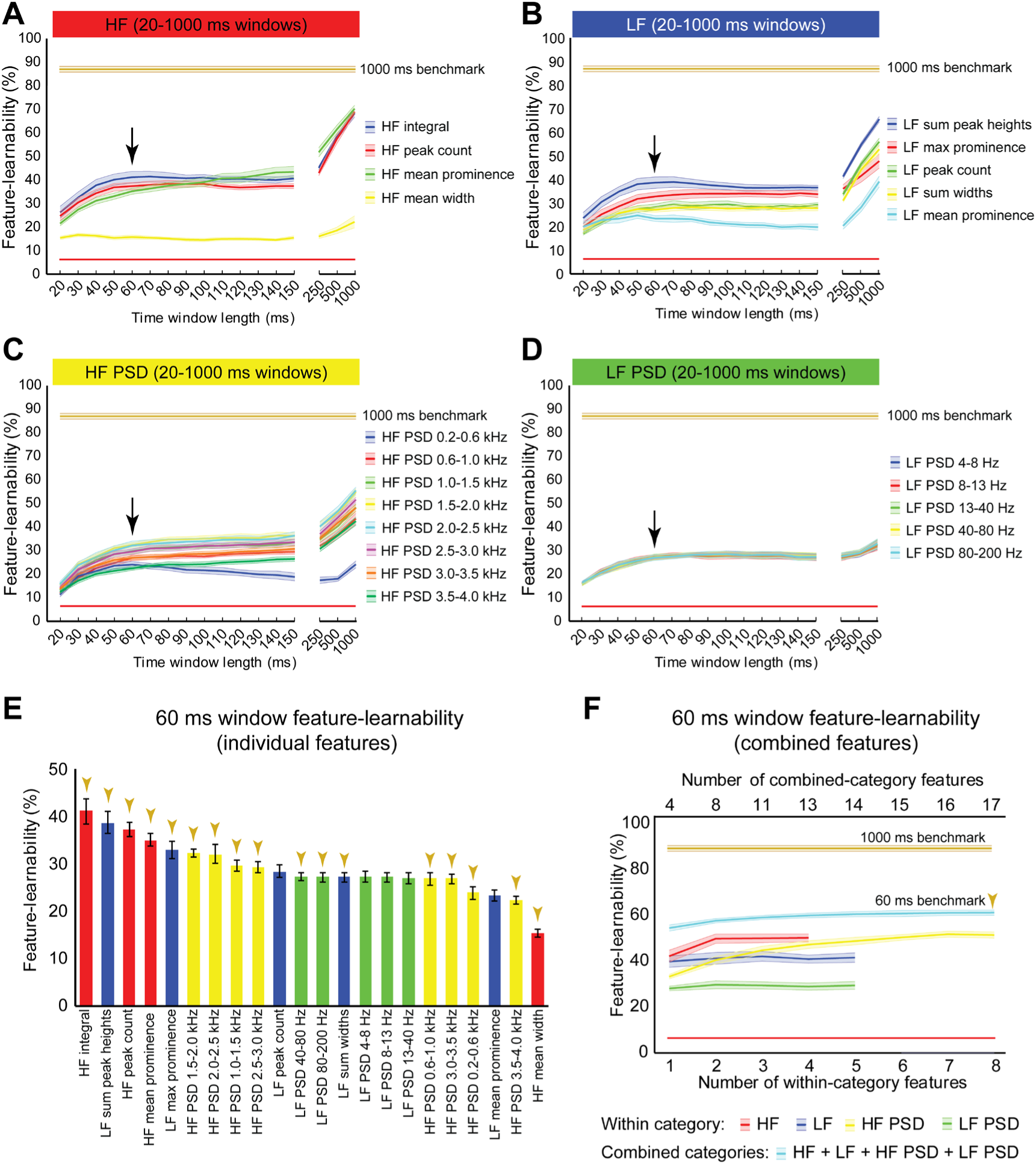
Benchmark signal features for an optimal short-time window. **(A-D)** Feature-learnability of features extracted from time windows of varying lengths from 20 ms to 1000 ms are shown in their HF **(A)**, LF **(B)**, HF PSD **(C)**, and LF PSD **(D)** signal feature categories. Coloured lines and shading indicate the feature-learnability (mean ± SEM) for each signal feature (colours arbitrarily chosen). Gold lines indicate the 1000 ms window benchmark feature-learnability ± SEM (see Figure 4D) for comparison; red straight lines indicate classification chance level (6.25%). Arrows indicate the 60 ms window as the optimised duration across all features where return of feature-learnability diminishes for increasing time windows. **(E)**: Feature-learnability ranking of individual signal features is shown for the optimised short-time window (i.e. 60 ms). Gold arrows indicate the 13 signal features that comprise the benchmark configuration. **(F)**: Benchmark signal features were determined for the optimised short-time window using the identical algorithm applied to determine the 1000 ms window benchmark feature set shown in Fig 3A; blue curve shows feature-learnability from combining across categories; gold arrow indicates benchmark configuration of 17 features (indicated in E). For comparison, the 1000 ms window benchmark feature-learnability is shown (gold curve). See Fig 2 for feature descriptions and abbreviations.

Combining the best two features (*HF integral* + *LF sum peak height*) did not significantly improve feature learnability (41.8 ± 2.2) compared to *HF integral* alone (41.0 ± 2.5, p = 0.09, paired *t*-test). A combination of 4 signal features that included the highest-ranked feature from each category extracted from 60 ms, significantly improved feature-learnability compared to most within-category combinations, with the exception of the combination of 2 or more HF features (p ≥ 0.76) and 5 or more HF PSD features (p ≥ 0.35, LMER, Tukey; Fig 6F). Also noteworthy was that feature-learnability from combining *HF integral* and *HF mean prominence* was significantly greater than all other within-category feature combinations, except for three or more combined HF PSD features, all of which were not significantly different (p ≥ 0.6, LMER, Tukey). The 60 ms benchmark feature set, determined by continual addition of the next best features from each category (where possible), resulted in a feature-learnability outcome of 59.4% from 17 features (gold arrows, Fig 6E-F).

### Temporal profiles of feature-learnability during mechanical stimuli

To measure how informative signal features, extracted over a short period, are throughout stimulus presentation, we determined a feature-learnability time-series by extracting DCN signal features from 60 ms rolling windows (Fig 7). We started by examining the feature-learnability temporal profile for the 60 ms benchmark 17-features set (indicated by gold arrows, Fig 6E). This produced a complex waveform (gold trace, Fig 7A) with three distinct peaks: the 1^st^ is an abrupt, relatively sharp peak coinciding with the stimulus onset; the 2^nd^ and 3^rd^ peaks were broadened, and peaked at the approximate midpoint of the stimuli and beginning of the stimulus-off/return phase, respectively. To establish which stimuli contributed to these peaks, neural networks were restricted to input/output data sets of tactile- or proprioception-dominated stimuli (Fig 7A, red and blue traces, respectively). This revealed that the 1^st^ peak arose from tactile-dominated stimuli, the 2^nd^ peak arose from proprioceptive-dominated stimuli, whereas the 3^rd^ peak arose from both tactile- and proprioceptive-dominated stimuli. Another interesting observation was that during the stimulus-off/return phase, feature-learnability was significantly elevated from chance levels, indicating that the input features during this period were informing the neural network of some information about the stimulation being performed.

**Fig 7.**
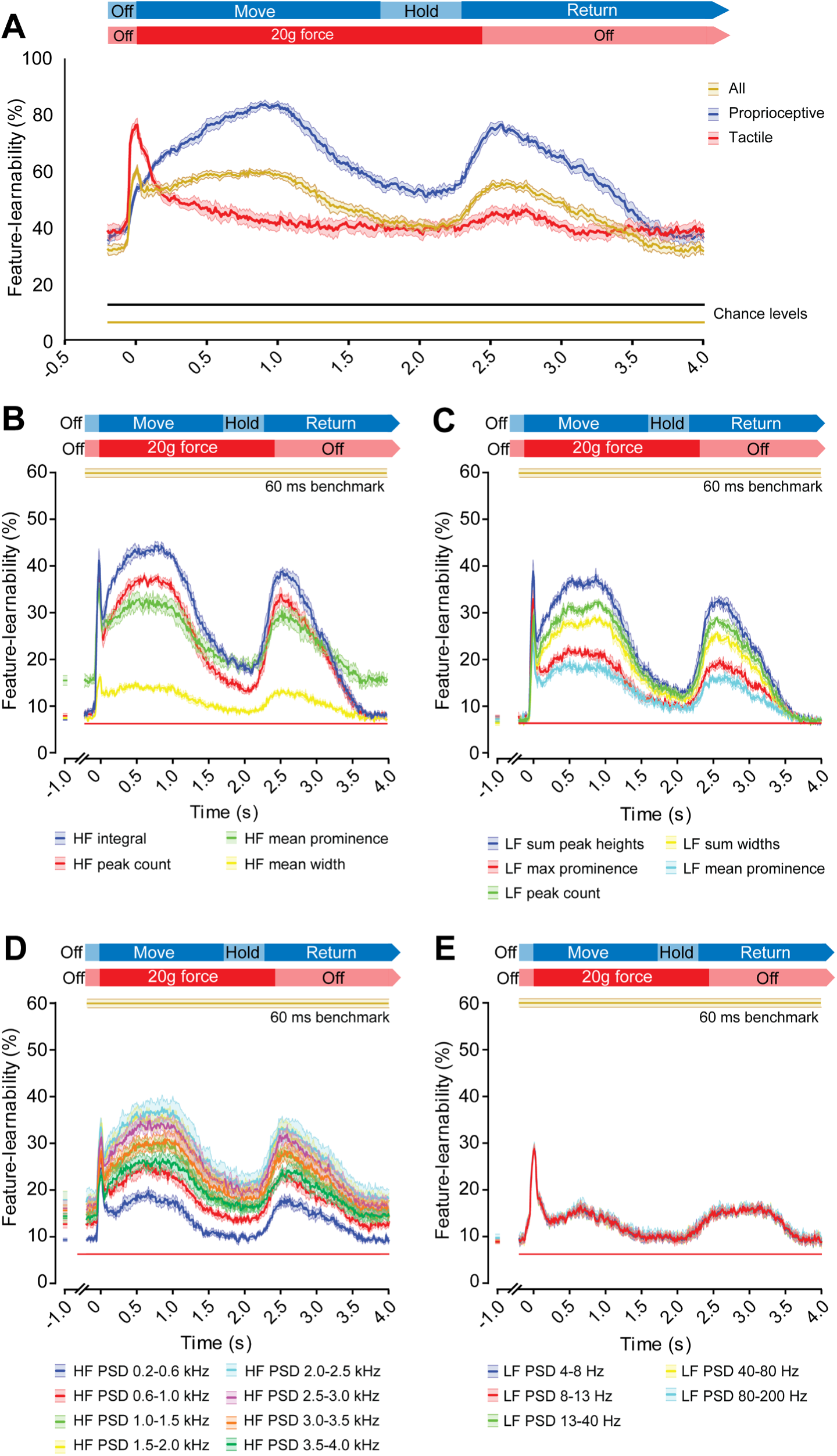
Evolution of feature-learnability from 60 ms windows during natural mechanical stimuli. **(A)** Feature-learnability time-series determined by extracting features from a 60 ms sliding window advanced every 10 ms are shown for the 60 ms benchmark feature set of 17 features (Fig 6E) for all stimuli (gold trace), as well as modified neural networks restricted to proprioceptive- or tactile-dominated inputs/outputs only (indicated by blue and red traces respectively). Expected feature-learnability by chance is indicated for all (gold) and proprioceptive- and tactile-dominated (black) stimuli. **(B-E)** Feature-learnability time-series at 1000 ms pre-stimulus and from 200 ms pre-stimulus to 4000 ms post-stimulus are shown in their HF **(B)**, LF **(C)**, HF PSD **(D)**, and LF PSD **(E)** signal feature categories. Coloured lines and shading indicate the feature-learnability (mean± SEM) for each signal feature (colours arbitrarily chosen). Gold lines indicate the 60 ms window benchmark feature-learnability ± SEM (Fig 6F) for comparison; red straight lines indicate classification chance level (6.25%). See Fig 2 for feature descriptions and abbreviations.

To determine how individual features contribute to feature-learnability over this time course, we examined feature-learnability time profiles for each of the 60 ms benchmark features (Fig 7B-D). For all features, the 3^rd^ peak coincided with the commencement of the off-stimulus period and their amplitudes never exceeded their respective 1^st^ or 2^nd^ peaks. Of the four time-domain HF features, the maximum peak for three of these coincided with the 1^st^ (tactile) peak, whereas the *HF integral* maximum fell on the 2^nd^ (proprioceptive) peak (Fig 7B). Feature-learnability returned to near-chance levels during the stimulus-off/return period for all HF features except *HF mean prominence*, which remained significantly elevated. Of the five time-domain LF features, the maximum peak for two of these coincided with the 1^st^ peak (*LF max prominence* and *LF mean prominence*), whereas the other three features had similar 1^st^ and 2^nd^ peak magnitudes (Fig 7C). Feature-learnability returned to chance levels during the stimulus-off/return period for all LF features.

Of the eight HF PSD features, the maximum peak coincided with the 1^st^ peak for two features derived from frequencies between 0.2-1.0 kHz, whereas maximum feature-learnability coincided with the 2^nd^ peak for features derived from frequencies between 1.5-4.0 kHz (Fig 7D). All HF PSD features remained elevated above chance levels during the stimulus-off period, with frequencies closer to the 2 kHz range demonstrating greater feature-learnability during the stimulus-off/return period. For all five LF PSD features, the maximum peak coincided with the 1^st^ peak, and all remained slightly elevated above chance levels during the stimulus-off/return period (Fig 7E).

To gain insight into why feature-learnability remained above chance levels during the stimulus-off/return period, we inspected confusion matrices from all animals for features associated with above-chance performance at 1 second pre-stimulus. Fig 8 shows examples of two features (*HF mean prominence* and *HF PSD 2.0-2.5 kHz*), which demonstrates that increased feature-learnability one second prior to stimulus arose for different reasons across different animals, but for similar reasons within animals. One common theme was that the correct limb was identified for the two stimulus categories. In all but one animal, one or more outputs were correctly classified with > 35% accuracy. To investigate if there were learning differences at the beginning vs the end of trials, we then divided data sets into thirds for i) each of the 10 trials, and ii) for the ~100 sequential stimulus presentations, to see if repeated stimuli within and across the trials contributed to improved outcomes. We found no difference in feature-learnability between the first-third and last-third of stimulus presentation within trial sets, or across all 100 trials, indicating that there were no changes to feature-learnability as a result of repeating trials.

**Fig 8.**
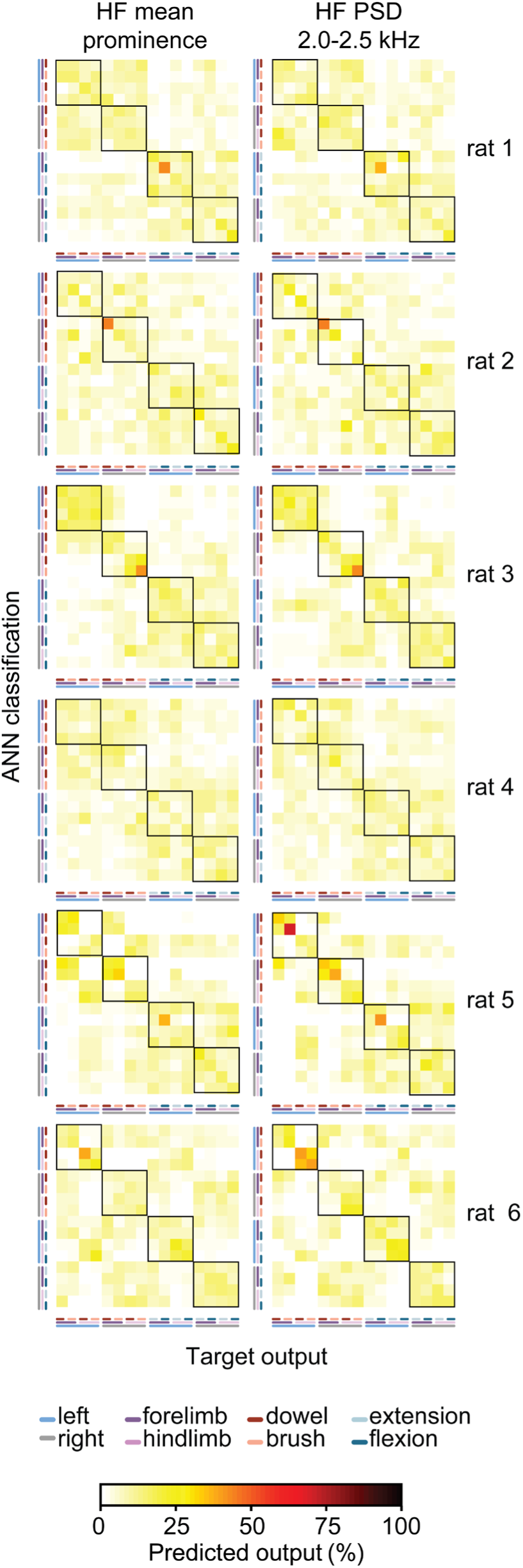
Individual animal machine learning outcomes for 60 ms window at 1 s pre-stimulus. Each confusion matrix shows the machine learning outcomes for individual animals (rows) for two features (columns) which demonstrated feature-learnability significantly above chance levels at 1 s pre-stimulus. Note the similarity within animals, but lack thereof across animals.

In summary, for 60 ms rolling windows some features performed better over periods rich in tactile-stimulus information and others over the proprioception-rich periods. PSD features derived from <1kHz, contributed more during tactile-rich periods, whereas those >1.5kHz contributed more toward proprioception-rich periods, while feature-learnability during the stimulus-off/return period was greatest for PSD features around 2 kHz and reduced for lower and higher frequencies. The reasons for above chance performance during the stimulus-off/return period appears non-generalisable because it was different for each animal.

## Discussion

Our study used feature-learnability to reveal several neural signal features that facilitate excellent decoding accuracy for mechanically-evoked tactile- and proprioception-dominated stimuli over a range of frequencies (4-4000 Hz). To our knowledge, some of these features and/or feature combinations have not previously been used to decode neural signals. We demonstrated that individual features extracted over 1000 ms of data from the HF category generally outperformed those from other categories, and that only two HF features from somatosensory DCN-signals were required to achieve the 1000 ms feature-learnability benchmark. With a shorter time-window of 60 ms – one that is compatible with neural prosthetic applications – decoding accuracy and robustness of signal features were greatly improved by adding relevant and diverse features, and reasonable classification accuracy was achieved despite sampling from electrodes with poor spatial resolution. We found proprioception-dominated stimuli were more accurately classified than tactile-dominated stimuli, and stimuli presented to the forelimbs were predicted better than hindlimbs. Our study established time courses that track how information relevance, for each feature, varies as a function of the mechanical stimulation phases. We discuss these findings below in relation to the underlying DCN physiology after the following methodological considerations.

The sMEA used to capture surface potentials had relatively poor spatial resolution and a low number of electrodes. Others have shown that recording with higher densities will permit greater classification accuracy (Mehring *et al.*, 2003; Bansal *et al.*, 2011; Wong *et al.*, 2016). Our sub-optimal sMEA arrangement is likely to limit classification accuracy compared to a sMEA with a higher number and density of electrodes, but this conservative electrode configuration allowed us to focus on how individual features contributed to decoding of somatosensory neural signals under less than ideal recording conditions. A more strategic recording electrode arrangement is likely to permit greater classification accuracy than our present findings and would be recommended for neural prosthetics applications. We and others have been previously identified the DCN as a potential neuroprosthetic target for centrally restoring somatosensation (Delhaye *et al.*, 2016; Richardson *et al.*, 2016; Loutit *et al.*, 2019). Our capacity to achieve high classification accuracy with a sub-optimal electrode configuration is supportive of the DCN neuroprosthetic potential because the ability to reproducibly represent the external environment encoded as neural signals in the DCN makes it amenable to artificially recreating signals using a biomimetic simulation approach (Saal & Bensmaia, 2015; George *et al.*, 2019).

We described our stimuli as either tactile- or proprioception-dominated. While these stimuli target most of the intended afferents under investigation, we must acknowledge the diversity of afferents being recruited by our natural stimuli. For example, the 20 g force applied by the tactile-dominated stimuli to the palmar/plantar surfaces are likely to have also moved wrist/ankle and finger/toe joints, while the proprioception-dominated stimuli are likely to have activated hair and skin afferents around joints, on resting surfaces of the limb, and the skin where the actuator was attached that moved the limbs. These stimuli therefore do not activate tactile and proprioception afferents in isolation; however, the time courses of neural responses were characteristic of tactile and proprioceptive afferent firing, indicating that the majority of intended afferents were appropriately activated. It is also worth noting that any natural stimulus will activate a mix of afferent populations and therefore activation of a pure afferent type would be a rare occurrence in nature.

### Prediction of somatosensory stimuli from dorsal column nuclei signals

Across all individual features and our benchmark feature set, proprioception-dominated stimuli were generally predicted better than tactile-dominated stimuli, and forelimbs were predicted better than hindlimbs. More accurate proprioception classification compared to tactile is likely to have resulted because there was a clear significant separation in the neural activity evoked by flexion and extension (indicated by the *HF peak count* feature alone), and both these proprioceptive stimuli evoked activity that was significantly different to both tactile stimuli, whereas, the significant difference and effect size was much less between the two tactile stimuli. Greater proprioception evoked activity may have resulted from activating more receptors and/or evoking more action potentials from each activated receptor. It is likely that the proprioception-dominated stimuli activated many more receptors over the entire limb, as hair, skin, muscles, tendons, and joints, were likely to be activated from the digits to the shoulder, whereas tactile-dominated stimuli were isolated to cutaneous receptors in the glabrous skin of the paws, and potentially activated some hairy skin on the back of the paws due to the downward force exerted. Furthermore, tactile stimuli evoked most activity at stimulus onset, then appeared to quickly adapt (Figs 1B & 7A), whereas moving the limb evoked high activity levels throughout the entire duration of both proprioceptive stimuli, thus the potentially higher number of receptors were also active for a longer duration.

Forelimbs may have been predicted better than hindlimbs because most hindlimb activity was acquired by the same midline electrodes which spanned across the two gracile nuclei on both sides, whereas each cuneate nucleus on either side had its own electrode, thereby facilitating better spatial discrimination. Some of the hindlimb errors resulted from confusing tactile- and proprioceptive-dominated stimuli of the same limb (Fig 4A-D and Fig 5B, E). Most hindlimb proprioceptive afferents either project onto DCN neurons in the ventral gracile nuclei or to nuclei X and Z, which are DCN accessory nuclei (Landgren & Silfvenius, 1971; Johansson & Silfvenius, 1977a, b; Mountcastle, 1984; Mantle‐St. John & Tracey, 1987). The proprioceptive DCN regions may be too deep to acquire HF features, while nuclei X and Z are not covered by the placement of our electrode array, thus, prediction of hindlimb stimuli may have relied on mostly tactile information to discriminate between the tactile and proprioceptive-dominated stimuli, which may also have contributed to lower hindlimb prediction accuracy.

### Feature-learnability within and across animals

To determine if benchmark features are generally useful for robust decoding of mechanical stimuli in different animals and not unique to individual animals, we quantified feature-learnability using the LOO approach. This approach trains the machine-learning algorithm on features extracted from all other animals and tests on the remaining animal. We previously established that comparing feature-learnability under LOO and WIA conditions provides insight into how well, or poorly, DCN-signal extracted features generalise across animals (Loutit *et al.*, 2019).

Forelimb flexion and extension were minimally perturbed under LOO conditions, indicating that 1000 ms benchmark features were highly relevant across all animals for forelimb proprioception stimuli. However, LOO conditions reduced feature-learnability by ~37%, which was mostly due to a more than 50% reduction in the ability to predict tactile stimuli and ~40% reduction in the ability to predict hindlimb proprioception-dominated stimuli. This indicates that a significant portion of information contained within benchmark features are no longer relevant, or as generalisable, for tactile- and hindlimb-presented stimuli across the different animals. It was evident from the WIA approach that, compared to proprioception, tactile stimuli were more challenging to classify, which arose almost exclusively from confusion errors between the dowel and brush stimuli rather than side and limb. The two tactile stimuli are challenging to discriminate because they present the same forces but differ only subtly by their textures on contact with the paw. The reduced classification accuracy under LOO conditions are likely to reflect a sufficiently large inter-animal variability of responses for the two stimuli; e.g., dowel responses in one animal produced features that appear similar to those induced by brush responses from another animal. Hindlimb stimuli classification errors (both tactile- and proprioception-dominated) mainly resulted from confusing the side of the body that a stimulus was presented. Signal asymmetry across the DCN surface, as we have previously shown for hindlimb stimulated nerves (Loutit *et al.*, 2019), could lead to larger signal variations in left and right limb-derived activity, across the different animals we observed in the present study. Furthermore, small left/right differences of midline electrode placement between animals could cause a non-generalisable variation in detected afferent activity across animals, and thereby contribute to reduced LOO hindlimb classification. Forelimb afferent activity is less sensitive to slight left/right variations in placement because the acquired afferent activity remains segregated on distinct electrodes. We therefore speculate that improvements to hindlimb decoding accuracy may be achieved by medially placed electrodes on either side of the midline.

Reducing thirteen features to two HF features revealed only minor differences under WIA and LOO conditions, indicating that when features are extracted over a 1000 ms period, the additional eleven features do not contribute substantial additional information toward decoding accuracy. When the feature extraction window was reduced to 60 ms under WIA conditions, combinations of the two highest-ranked, or two best HF features, no longer reached the feature-learnability obtained by the 17-feature 60 ms benchmark, indicating that the additional fifteen features did contribute additional information over the shorter time period. However, as the 17-feature 60 ms benchmark did not reach the feature-learnability obtained by the two best HF features extracted over 1000 ms, the information contained by the two HF features over that additional period (i.e. 940 ms) was more valuable than all benchmark features extracted over the 60 ms. This demonstrates that reducing time windows of feature extraction reduces information content, in which case adding other relevant features provides some compensation in feature-learnability performance.

### High-frequency features

*HF mean prominence* was the best performing feature for the 1000 ms windows, and fourth in the 60 ms window, yet, to our knowledge this signal feature has not been previously used for neural decoding. The *HF mean prominence* feature resulted in fewer errors when classifying brush and dowel stimuli of the same limb, compared to all other features (Fig 4), which implies that the machine-learning algorithm detected greater contrast in this feature’s magnitude under the two tactile stimulus conditions compared to other features. We observed that at stimulus onset, dowel stimuli produced more precisely timed bursts of activity than brush stimuli. This may have facilitated the improved dowel/brush classification from the *HF mean prominence* feature because spikes from multiple neurons or afferent fibres arriving at an electrode simultaneously (i.e. responses evoked by dowel stimuli) will summate, and therefore show higher average peak prominences than single spikes that have less temporal overlap (i.e. brush stimulus evoked responses). This explanation is supported by the observed significantly greater *HF mean prominence* evoked by dowel, compared to brush stimuli.

The combination of *HF mean prominence* and *HF peak count* features, extracted over 1000 ms, was sufficient to achieve feature-learnability performance equivalent to the 87% feature-learnability benchmark. Furthermore, this 2-feature input configuration resulted in no confusion errors under WIA, and very few under LOO conditions, between the tactile and proprioception stimulus categories (i.e. no and few confusion errors in lower left and upper right quadrants of Fig 5D and E respectively). This result can be explained by the significant difference for *HF mean prominence* and the > 2.5-fold difference in *HF peak count* when tactile-, compared to proprioception-dominated stimuli were used.

For a fixed time-window, *HF peak count* provides information about spike frequency and the total number of action potentials generated by peripheral afferents. The large difference in this feature between tactile- and proprioception-dominated stimuli may have resulted from the relatively small vs. larger receptive field sizes respectively that these stimuli engaged. *HF mean prominence* captures spiking temporal alignment (as discussed above) and spike magnitudes. Spike magnitudes are influence by neuron and or axon size, as well as the event’s distance from the electrode (Gasser & Grundfest, 1939; Hunt, 1951; Nelson, 1966; Buchwald & Grover, 1970; Grover & Buchwald, 1970). Group 1 proprioceptive afferents generally have larger diameters than Aβ tactile afferents(Gasser, 1941; Kandel *et al.*, 2000). In addition to the laterally placed external cuneate nucleus, proprioceptive afferents preferentially terminate deep on ventral DCN neurons (Campbell *et al.*, 1974), which have a high proportion of large somas (Cheema *et al.*, 1983) that are located 500-700 µm below the brainstem surface in rats (Li *et al.*, 2012). Tactile afferents terminate on smaller neurons in a cluster zone that are located at approximately half the depth as the ventral DCN neurons (Li *et al.*, 2012). How these two neuronal populations (larger somas located deeper vs smaller somas located more superficially) affect spike magnitudes measured from the surface are difficult to assess without dedicated experiments, however, it is likely that these anatomically segregated neuronal populations evoke different shaped spike waveforms. The combination of stimulus evoked spike-timing, the signal location and neuronal types responsible for generating the signal may therefore result in unique prominence profiles for each of the stimuli, and thereby contribute to improved stimulus classification.

### Low-frequency features

When rectified HF signals were low-pass filtered, peaks in the LF signals occurred over periods with high rates of HF events, separated by troughs with few or no HF events. Therefore, the LF features may reveal information about bursting properties of a neural population. The LF feature category demonstrated varied feature-learnability, with its best performer, *LF sum peak heights*, ranking in the top four features in the 1000 ms and 60 ms windows. This feature likely represents very similar activity to *HF integral*, as both are extracted from rectified HF signals, and summing envelope peaks captures related information to the integral of the signal. Congruently, *LF sum peak heights* and *HF integral* showed very similar feature-learnability characteristics (compare Figs 3A, 6E, 7B and 7C) and confusion errors (Fig 4A-B), and combining these two features did not significantly improve feature-learnability from *HF integral* alone. These features may capture a combination of HF events, slower synaptic activity, or subthreshold events, representing a measure of the signal energy in a neural population.

### Power spectral density features

The HF PSD feature set was similar to that used by Bouton *et al.* (2016) who extracted features over 100 ms windows for motor signal decoding in a brain-machine interface capable of effecting limb movement through neuromuscular stimulation. This feature set represents frequency power in the multiunit activity range, which is typically considered to be about 300-6000 Hz (Stark & Abeles, 2007) (our range: 200-4000 Hz). We found these features generally to be good predictors of somatosensory DCN signals. In our 60 ms window data, the frequency bands of 1.5-2.5 kHz were of interest because 1) the highest decoding for both tactile- and proprioception-dominated stimuli were found for this feature category over this range, and 2) frequency bands above 1.5 kHz showed a peak in the proprioception-dominated phase, while bands below 1.5 kHz, including the LF PSD features, had peaks in the tactile-dominated phase. Each of the eight HF PSD bands appeared to add some unique information not captured by the other bands, as successive additions of these features continued to improve feature-learnability until all the HF PSD features were exhausted (see yellow line, Fig 6F). However, feature-learnability derived from all eight HF bands was not significantly greater than combining *HF integral* and *HF mean prominence*, indicating that these features could offer superior neural decoding compared to all HF PSD features.

### Feature-learnability temporal profiles

Tactile-dominated stimuli evoked small bursts of activity at stimulus onset and offset, while proprioception-dominated stimuli evoked little activity at stimulus onset, but larger amounts during the middle of the stimulus and return periods on anatomically relevant electrodes (Fig 1B). These phases of activity were captured by the peaks of the feature-learnability temporal profile (gold curve, Fig. 7A). We can confirm that the 1^st^ peak of this temporal profile reflects the period where considerable tactile information is presented to the artificial neural network because this peak is abolished when tactile data were omitted from the learning algorithm (blue curve, Fig 7A), but remains when tactile data is present and proprioceptive data were omitted (red curve, Fig 7A). Conversely, we can confirm that the 2^nd^ peak is attributed to the duration when maximum proprioceptive information is present. The feature-learnability temporal profile of individual features can therefore provide insight into their capacity to provide decoding information throughout the progression of the stimuli. For example, it is evident that LF PSD signal features below 200 Hz contain relatively little information for decoding proprioceptive stimuli, whereas they contribute significantly toward decoding tactile events. Interestingly, the tactile-dominated feature-learnability temporal profile appears to reflect stereotyped afferent activity adaptation characteristics, whereas the proprioception feature-learnability temporal profile appears to reflect the dynamic movement of the proprioceptive stimuli, with its peak coinciding when the limbs were maximally flexed or extended (i.e. at the midpoint of the *Move* phase, Fig 7).

### Rest period activity

Feature learnability remained above chance levels 1 second before stimuli were presented (Fig 7) for the *HF mean prominence* and all HF PSD features. This indicates that during the rest periods, some information specific to the limb and/or the stimulus being presented was encoded in these DCN signal features. That the more successful predictions were not consistent across all animals (Fig 8), suggests that the prediction is not based on some residual activity from particular stimuli, but rather, multiple features in the same animal contributed information that was unique to that animal for a combination of stimulus and location. Furthermore, it was unlikely that above-chance level classification arose from some learned effect during repetitive stimulus presentations because feature-learnability was not different when comparing the first and last third of trials, although, it should be noted that training/testing data sets were also reduced to one third, making accurate classification more challenging.

What then could account for this predictive capacity prior to the stimuli? One possibility is that the limb undergoing repetitive stimulation experienced a different afferent activation status during the stimulus-off period compared to the remaining limbs. This could have arisen from continued reactivation of slowly adapting afferent activity during the off-stimulus period because of the continued interruption of the repetitive stimulation cycles, whereas the slowly adapting activity from the non-stimulated limbs would have greater opportunity to cease firing. An alternative explanation could be from activity resulting from long-lasting membrane potential depolarisations seen in DCN cells in response to sensory stimulation (Canedo *et al.*, 1998), or rhythmic activity that outlasts stimulation periods (Nuñez & Buño, 1999). We have not found evidence from other studies finding similarly high decoding accuracy in stimulus off periods. Nevertheless, this phenomenon demonstrates remarkable sensitivity of the HF PSD and *HF mean prominence* features for capturing status differences in afferent populations.

## Conclusion

Feature-learnability enables us to assess the information contained in DCN surface potentials for decoding natural tactile- and proprioceptive-dominated somatosensory events. We identified individual, and combinations of signal features with superior decoding capacity, some of which, to our knowledge, are not currently routinely used for neural decoding. Generally, HF time-domain features are most informative for decoding somatosensory-evoked neural signals compared to frequency-domain features, which is likely to translate to the decoding of motor neural signals. For sufficiently large time windows, only two HF time-domain features are adequate to achieve benchmark decoding accuracy, but for shorter time windows that are more practical for neural prosthetic applications, increasing the number and diversity of features improves the decoding robustness and accuracy. We showed that a feature’s decoding capacity is altered throughout a dynamic event. Future neural decoding algorithms may select features for their capacity to specialise in contributing unique information at different points in time during complex sensorimotor tasks. The signal features and evaluation approach we use here may improve neural decoding for machine-learning classification applications such as future neural prosthesis design. The high decoding accuracy achieved in this study is supportive of the DCN as a possible future somatosensory neural prosthetic target.

## Materials and Methods

### Animals

We used 8-week-old male Wistar rats (283-464g; n = 6) (Australian Phenomics Facility, Canberra, ACT, Australia). There were 1-3 animals housed per cage, with a 12/12-hour light/dark cycle. Food and water were available for animals to access *ad libitum*. All procedures were approved by the Australian National University Animal Experimentation Ethics Committee (A2014/52) and adhered to the Australian code of practice for the care and use of animals for scientific purposes.

### Surgery

Rats were anaesthetised with urethane (1.4g kg^−1^ i.p.). A tracheotomy was performed, and a breathing tube inserted to aid natural respiration. Rats were placed in a stereotaxic frame with head flexion at −20⁰ (Stoelting Instruments). The dorsal skin and muscles of the neck, and the dura and arachnoid mater were excised between the foramen magnum and the C1 vertebra to expose the dorsal surface of the brainstem. In most cases the C1 vertebra was cut away with rongeurs to aid placement of an adapted surface multi-electrode array (sMEA; *Nucleus 22 Auditory Brainstem Implant*, Cochlear Ltd.; see Chelvanayagam *et al.* (2008)for details). A flexible plastic rod held by a micromanipulator on the stereotaxic frame was lightly pressed onto the sMEA to hold it symmetrically over the brainstem midline.

### Stimulation and recording

We applied two types each of tactile- and proprioception-dominated stimuli to the rat limbs amounting to sixteen (4 stimuli × 4 limbs) possible mechanical stimulus conditions. The tactile-dominated stimuli were applied by pressing either a wooden dowel rod (diameter: 2.0 mm) or brush (tip diameter: 1.0 mm) into the palmar/plantar pads of the paws. The rod and brush were fixed at the end of a flexible tube to deliver 20 g of force. The proprioception-dominated stimuli were applied to the rat by flexing or extending the rat’s limbs. The rats’ limbs were fixed to a dowel rod with adhesive (Loctite) which enabled the experimenter to directly manipulate the limbs. All stimuli were applied by the experimenter who was triggered by the timing of a metronome.

Stimuli were applied for 2.4 seconds with a rest period of 2.4 seconds. Tactile-dominated stimuli had a stimulus on/off pattern (4.8 seconds per trial), but proprioception-dominated stimuli also had a 2.4 second period which was used to return the limb to the resting state, where it remained in resting state for 2.4 seconds (7.2 seconds per trial). Stimuli were applied in 10 sets of ~10 trials (~100 stimuli per type), with ~30 seconds rest between sets.

DCN electrical signals were acquired through seven electrodes of the sMEA and filtered (50 Hz notch filter; 10 kHz low-pass filter) through custom built amplifiers. The signals were then digitally recorded (40 kHz sample rate) through a PowerLab 16/35 acquisition system and viewed in LabChart Pro software (Version 8.1.1, ADInstruments, Bella Vista, NSW, Australia). The seven electrodes of the sMEA are referred to from rostral to caudal as follows: left side electrodes: e1, e2; midline electrodes: e3, e4, e5; right side electrodes: e6, e7 (Fig 1A, right insert), and all seven electrodes combined are referred to as e{1-7}.

### Signal processing and feature extraction

Signal processing, feature extraction, and analysis were performed offline (MATLAB version R2018a, MathWorks). We used twenty-two DCN signal features as artificial neural network inputs. The names, description, and examples of feature extraction for all twenty-two features are shown in Fig 2. Four features were extracted from high-frequency filtered (bandpass 0.55-3.3 kHZ; 5-order Butterworth filter) signals (HF features; Fig 2A-D), and five from low-frequency signals (LF features; Fig 2E-I), which were quantified from the HF signals after they had been rectified and low-pass filtered (< 80 Hz; 5-order Butterworth filter). Thirteen features were extracted from frequency spectrograms (*spectrogram* MATLAB function) of DCN signals bandpass filtered between 4-5000 Hz (8-order Butterworth). Eight of these features were quantified from the peak power spectral density of high-frequency bands (HF PSD features): 200-600 Hz, 600-1000 Hz, 1000-1500 Hz, 1500-2000 Hz, 2000-2500 Hz, 2500-3000 Hz, 3000-3500 Hz, 3500-4000 Hz (Fig 2J). The other five spectral features were quantified from the peak power spectral density of low-frequency bands (LF PSD features): 4-8 Hz, 8-13 Hz, 13-40 Hz, 40-80 Hz, 80-200 Hz (Fig 2K). For all filtering we used a zero-phase response filter: *filtfilt* function (MATLAB).

### Standardised artificial neural network for machine-learning

Individual signal features extracted from each of the seven electrodes were paired to the stimuli that generated them, to create input/output pairs for machine-learning. All machine-learning experiments used a standardised artificial neural network (ANN) with a supervised learning classification algorithm (*patternnet* MATLAB function). The standardised ANN comprised 42 hidden neurons and 16 output neurons corresponding to the 16 possible stimuli (4 stimulus types, presented to 4 different limbs). We selected the number of hidden neurons by increasing their number between 16 (the number of outputs) and 154 (the highest possible number of inputs from our feature set). For each number of hidden neurons, we determined the average feature-learnability with the minimum (7) and maximum (154) number of features included in the input set and found that 42 hidden neurons produced the highest average feature-learnability. Thus, only the input neurons were altered, depending on the number of input features required for each experiment. The ANN used gradient descent with momentum and the adaptive learning rate backpropagation training function *traingdx*. The hyperbolic tangent sigmoid function, *tansig*, was used in the hidden layer units, and a softmax transfer function (*softmax*) was used in the output layer. Both inputs and output targets were normalized such that all values fit between −1 and 1, as per the *patternnet* default setting. For training, cross-validation, and testing, the inputs were separated into training, validation, and test subsets, using the *patternnet* default settings.

### Feature-learnability of individual features and selection of benchmark input feature sets

Feature-learnability provides a stable measure of relevant information content provided by input features including a measure of the information content variability (Loutit *et al.*, 2019). We used the Within Individual Animal (WIA) approach (Loutit *et al.*, 2019) for all feature-learnability testing, unless otherwise specified. In this approach, input/output pairs from ~1600 stimuli were generated for each individual animal. For each animal, ten repeated training (70%), validation (15%), and testing (15%) machine-learning cycles were performed under random initialising conditions. These ten confusion matrices were averaged to produce a single representative confusion matrix for each animal, then the representative confusion matrices for all animals were averaged. Feature-learnability was determined by the mean ± SEM derived from the diagonal of this matrix. To determine the feature-learnability of an individual feature, the inputs to the standardised ANN were restricted to the single input feature in question, resulting in a total of 7 inputs (the signal feature extracted from each electrode).

To obtain a measure that represents the maximum possible information content contained in signal features, a benchmark feature set was determined that produces the highest feature-learnability outcomes. This feature-learnability benchmark facilitates comparisons of information content contained by individual or subsets of input features. Individual features were extracted from all 7 electrodes over the first 1000 ms from stimulus onset, and feature-learnability determined. Individual features were then ranked from highest to lowest feature-learnability, prioritised by the largest means and smallest SEM (Loutit *et al.*, 2019). Input feature selection was performed using an adapted sequential forward floating search algorithm (Whitney, 1971; Pudil *et al.*, 1994). For the adapted search algorithm, we first tested feature-learnability improvements after including new features, from within the same feature category, to the input set (see Fig 2 for categories). Once we established the order that features were added to the feature set for each category, we used the within-category order for a second across-category search algorithm. The second search included the best four features per round – one feature from each category, unless all features from a category had been exhausted, in which case less than four features were added – and feature-learnability was recorded in each round of +4 feature additions. After each round of +4 additions, we also tested feature-learnability after removing each feature from the current feature set, in a backward search. If feature-learnability was improved by removing a feature from the feature-set, then that feature was kept out of the feature set. If more than one feature’s removal improved feature-learnability, then whichever feature’s removal resulted in the highest feature-learnability was kept out of the feature set. This removal process was repeated until feature-learnability no longer improved, after which the forward search was resumed. From these sequential rounds, we determined the feature set that was no longer improved by subsequent feature additions and subtractions, and deemed it the benchmark feature set.

A second benchmark was determined for features extracted over an optimal short-time window length (described below). The identical approach was used as for the 1000 ms window benchmark, except that features were extracted from the shorter time course.

### Signal feature robustness across animals

To determine the generalisability of the features across animals we used the 1000 ms window benchmark feature set to compare feature-learnability in two across-animal approaches. For the leave-one-out (LOO) approach, we randomly assigned input/output pairs from 5 animals into training (70%) and validation (30%) sets, and testing was performed on the remaining animal (100%). The LOO approach was applied such that each of the six animals was examined as the test data set once. For more details on these methods see (Loutit *et al.*, 2019).

### Feature-learnability of different time windows and optimal short-time window selection

We sought to determine feature-learnability based on a time window that would be compatible with neural prosthetic devices that could rapidly update peripheral sensory status information to the central nervous system at any given time. Typical motor decoding algorithms use time windows ranging from 50-100 ms (Lebedev & Nicolelis, 2017), which trades classification accuracy for a feasible reduction in time required for neural prosthetic applications.

To determine an optimal short-time window that provides the highest feature-learnability in the shortest time, we tested windows between 20 ms to 150 ms with 10 ms increments, 250 ms, and 500 ms, in addition to the 1000 ms windows described above. All windows started at the time of stimulus onset. We plotted feature-learnability against time windows for each feature and used the *findchangepts* MATLAB function to determine the most abrupt change in feature-learnability. We then averaged the changepoint values across all features and rounded to the nearest time window. A benchmark set of signal features was determined for the optimal short-time window (described above).

### Feature-learnability variations in time during stimulus presentation

Once we established the optimal short-time window, we sought to use the short rolling window to extract features throughout the duration of stimulus presentation and determine how feature-learnability varied over time. To do this we extracted the twenty-two features from the optimal short-time window, sampled at 1000 ms prior to the stimulus, then every 10 ms from 200 ms pre-stimulus, to 4000 ms post-stimulus. Finally, we tested feature-learnability for each of the features extracted from each of the 421 windows.

### Statistical analysis

We used R (version 3.4.4) (R Core Team, 2018) for all statistical analyses with the RStudio integrated development environment (version 1.1.442). For comparison of the feature-learnability benchmark within and across animals we used a linear model (LM; R *lm* function, *stats* package). For all other comparisons we used a linear mixed effects model (LMER; *lmer* function from the *lmerTest* package, (Kuznetzova *et al.*, 2019)). Multiple comparisons were performed using estimated marginal means comparisons with Tukey p-value adjustments (R *emmeans* function, *emmeans* package (Lenth *et al.*, 2019)). We verified model fitting by examining model residuals for normality. Where required, *log* or *logit* transformations (*car* package, (Fox *et al.*, 2019)) were applied to the data prior to statistical modelling. All data is expressed as means ± SEM unless otherwise stated. Probabilities of *p* < 0.05 were deemed significant.

## Acknowledgements

The authors are extremely grateful to the Bootes Medical Research Foundation which funded this project.

## Competing interests

The authors declare no competing interests.

